# Spatio-temporal analysis of human preimplantation development reveals dynamics of epiblast and trophectoderm

**DOI:** 10.1101/604751

**Authors:** Dimitri Meistermann, Sophie Loubersac, Arnaud Reignier, Julie Firmin, Valentin Francois Campion, Stéphanie Kilens, Yohann Lelièvre, Jenna Lammers, Magalie Feyeux, Phillipe Hulin, Steven Nedellec, Betty Bretin, Simon Covin, Gael Castel, Audrey Bihouée, Magali Soumillon, Tarjei Mikkelsen, Paul Barrière, Jérémie Bourdon, Thomas Fréour, Laurent David

## Abstract

Recent technological advances such as single-cell RNAseq^1-3^ and CRISPR-CAS9-mediated knock-out^4^ have allowed an unprecedented access into processes orchestrating human preimplantation development^5^. However, the sequence of events which occur during human preimplantation development are still unknown. In particular, timing of first human lineage specification, the process by which the morula cells acquire a specific fate, remains elusive. Here, we present a human preimplantation development model based on transcriptomic pseudotime modelling of scRNAseq biologically validated by spatial information and precise time-lapse staging. In contrast to mouse, we show that trophectoderm (TE) / inner cell mass (ICM) lineage specification in human is only detectable at the transcriptomic level at the blastocyst stage, just prior to expansion. We validated the expression profile of novel markers enabling precise staging of human preimplantation embryos, such as IFI16 which highlights establishment of epiblast (EPI) and NR2F2 which appears at the transition from specified to mature TE. Strikingly, mature TE cells arise from the polar side, just after specification, supporting a model of polar TE cells driving TE maturation. Altogether, our study unravels the first lineage specification event in the human embryo and provides a browsable resource for mapping spatio-temporal events underlying human lineage specification.

---

Mammalian lineage specification has mostly been studied in mouse, using a combination of genetic manipulations, lineage tracking and immunofluorescence, leading to two-steps model of lineage specification. The first specification, segregating TE from ICM, occurs at the morula stage and is driven by a YAP/TEAD4/CDX2 axis^5, 6^. The second specification, separating ICM cells into EPI and primitive endoderm (PE), occurs at the blastocyst stage, which is driven by a NANOG/FGF4/GATA6 axis^5, 6^. scRNAseq offers the opportunity to understand the continuum between unspecified and specified cells^7^. To do so, we computed a pseudotime model with reversed-graph embedded projection to generate cell fate trajectories: the likeliest path that temporally orders cells based on their transcriptome^8^. This mouse pseudotime model shows transcriptomic changes immediately at the late 16 stage, with TE/ICM specification achieved by 32 cell stage, in line with our current understanding of mouse preimplantation development^9^ (**Extended Data Fig. 1a-f**).

A problem impairing the analysis of scRNA-seq datasets of human embryos is the asynchrony of *in vitro* culture. For *in vitro* fertilization (IVF) cycles, morphology-based staging are preferred to time, especially as multiple morphological events occur during embryonic day 5 (E5)^10^ (**Extended Data Fig. 1g**). Time-lapse analysis coupled with the pseudotime allowed us to link morphology-based stage and transcriptomic events. Embryos were laser-dissected to separate the mural TE from the polar TE/ICM (**Extended Data Fig. 2a-c**). Our set of 24 embryos and 150 cells were pooled with available datasets^1, 2, 4^, totaling 128 embryos and 1751 cells. Combining our in-depth annotated dataset with available datasets gave us the opportunity to generate a biologically validated model. Normalization and batch correction were applied^11^ and a pseudotime model was generated, with all 4 datasets interspersed (**Extended Data Fig. 2d-f**). The resulting human pseudotime model is characterized by 5 branches with 2 branching points; pseudotime value indicated the transcriptomic distance from the root cell (**Fig. 1a**). Projection of cells based on their developmental stage revealed the 8-cell stage to be the root, followed by the morula and blastocysts; projection of cells based on their dissection annotation showed mural TE cells in one branch (**Fig. 1b**). Branch identification was confirmed by enrichment in specific branches of human markers validated for EPI (KLF17), PE (SOX17) and TE (GATA2) (**Fig. 1c**, **Extended Data Fig. 3a**)^12^.

**Figure 1.**
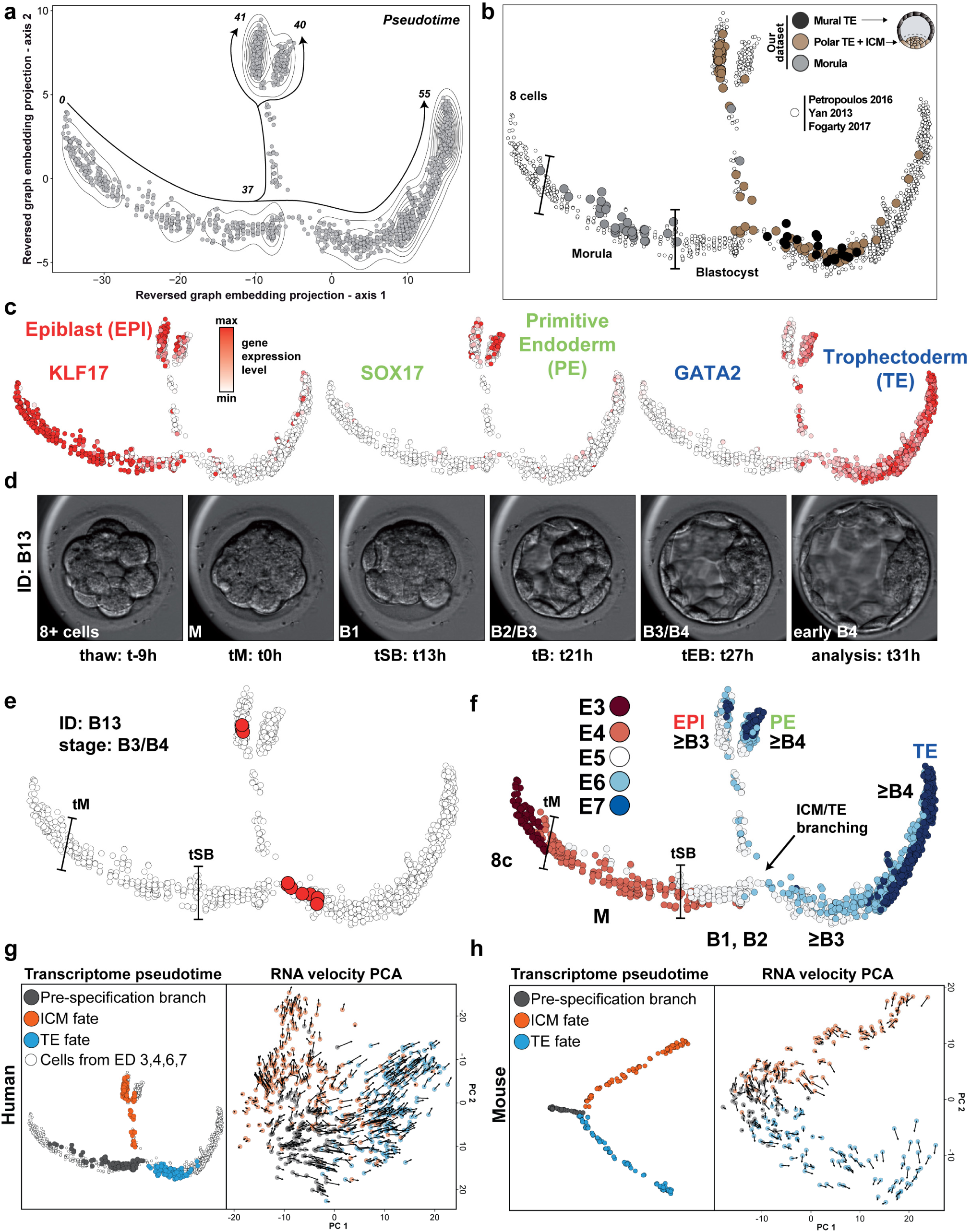
Transcriptomic model of Human preimplantation embryo development. **a**, Projection of scRNAseq samples with reversed graph embedding method (Monocle2). Density curves indicate sample density; pseudotime values represent transcriptomic distance from the root cell. **b**, Projection of our sample’s annotation on the pseudotime model. Morula cells (grey) and blastocyst dissection origin of cells are indicated (mural trophectoderm: black, polar trophectoderm or inner cell mass: brown). **c**, Projection of lineage marker expression levels: KLF17 (epiblast), SOX17 (primitive endoderm), and GATA2 (trophectoderm). **d-e**, Frames from time-lapse microscopy for embryo “B13”. For each embryo sequenced in this study, their morphokinetics were acquired by time-lapse microscopy (**d**). Developmental events include: morula compaction (tM), blastulation (tSB) leading to B1 stage, full blastocyst (tB) at B3 stage and expanded blastocyst once the zona pelucida thickness is halved (tEB) at B4 stage. tM is used as t0 to compare thawed embryos. A projection of cells from embryo B13 on the pseudotime model shows the correspondence between the pseudotime and the stage of the embryo (**e**). **f**, Projection of developmental day (E3 to E7) for all samples combined for this study, and the result of our refined staging. **g-h**, Transcriptome pseudotime and RNA velocity comparison of cells before and after specification in human (**g**) and mouse (**h**). For human, only embryos at day 5 were used in the RNA velocity principal component analysis.

To link developmental stage and molecular events, we projected all cells from each embryo on the pseudotime maps. For example, projection of blastocyst ID=13 (B4 stage upon scRNAseq analysis) (**Fig. 1d**) showed that at this stage, some cells are populating the EPI branch while other cells are populating the beginning of the TE branch, confirming that specification has occurred at this stage (**Fig. 1e**). Our annotation based on time-lapse and zona pellucida thickness on all our embryos shows that the first lineage specification in human occurs between the B2 and B3 stages, after the beginning of blastocyst cavitation (**Extended Data Fig. 4** and **Fig. 1f**). This resolves annotation overlap between E4, E5 and E6 in previous dataset, which limited their translation into sequence of events pacing human preimplantation development (**Extended Data Fig. 5**). Indeed, during E5, human embryos progress from early blastocysts (B1 and B2) to blastocysts (B3) and expanded blastocysts (B4). Finally, we projected cells from OCT4 KO human embryos^4^. This showed that OCT4 KO cells passed the ICM/TE specification point, but failed to reach mature EPI or TE branches, highlighting how our model can be used to refine human preimplantation studies (**Extended Data Fig.6a**). We developed an online tool to browse gene expression and cell annotation, providing a resource stimulating research efforts of human preimplantation development (**Extended Data Fig.6b-c**).

A puzzling observation is that in contrasts to mouse, positional information in human is not coupled to transcriptomic changes at the morula stage (**Fig. 1f**). Recently developed computational analyses allow interrogation of RNA velocity, a measurement of non-spliced mRNA^13^. This strategy reveals mRNA being actively transcribed in cells and not yet translated, thereby indicating cellular fate. RNA velocity analysis of cells before and after specification in mouse and human revealed cells with similar transcriptomic signatures, but with velocity trending toward ICM or TE, supporting our pseudotime models (**Fig. 1g-h**). Those observations are in line with the fact that cells at B2 stage co-express KLF17 and GATA2 at the mRNA and protein level, but are enriched in either KLF17 or GATA2 RNA velocity (**Extended Data Fig. 3b**). Altogether, transcriptomic modeling of mouse and human lineage specification shows a remarkable difference in the timing between species and identified the first lineage specification at B2/B3 stage in human, whereas it occurs at the morula stage in mouse.

To further investigate whether multiple states were present within each branch, we performed a gene-based analysis. 8 major modules of co-expressed genes were identified using weighted correlation network analysis (WGCNA)^14^, subdividing the pseudotime into 9 states defined by absence or presence of gene module (**Fig. 2a**). Modules are named after one of its representative gene. Each module contains specific expression dynamics, linked to specific transcription factors and enriched functions (**Extended Data Fig. 7 & 8**). Some genes modules reflected overall changes in the embryos: ZSCAN4 and DUXA modules are associated with zygotic genome activation between 8-cell and morula stages; the DNMT3L module is emerging in all cells at the blastocyst stage, and contains epigenetic regulators in line with methylation dynamics in human preimplantation development^15^ and metabolic pathways. The POU5F1B module contains pluripotency genes. Its expression starts in all cells of the morula, and then is restricted to the mature epiblast. This make POU5F1B module the first expressed module that has a cell lineage specific expression later on. TE cells from early blastocysts (B3) are plastic and can give rise to ICM cells but this plasticity is lost in expanded blastocysts (B4)^16^, which correlates with loss of expression of POU5F1B module in the TE. Other modules are related to specified lineages: IFI16 module for mature/stable EPI, GATA4 module for PE, GATA2 module for early TE and TE, and NR2F2 module for mature TE.

**Figure 2.**
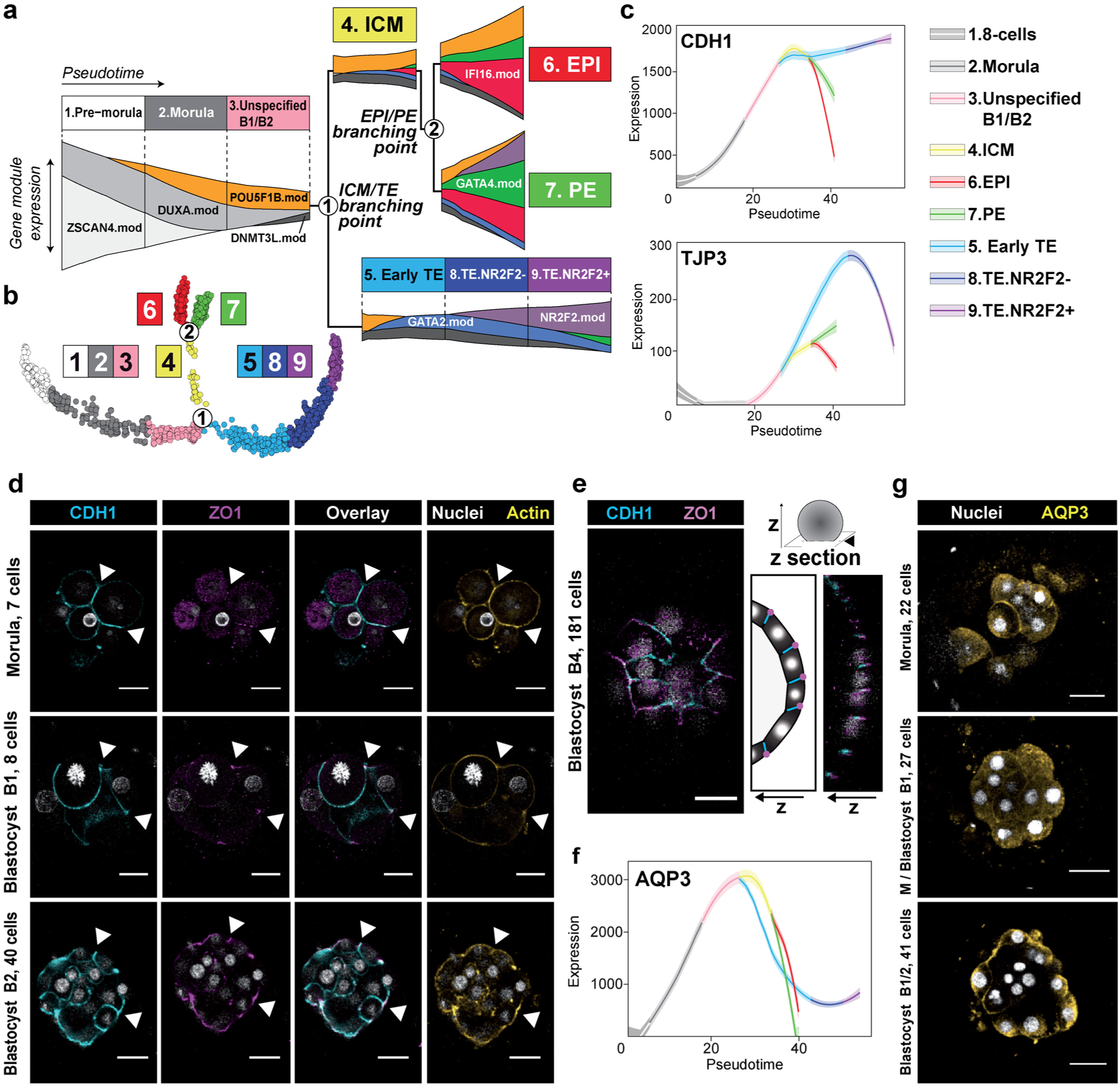
Defining spatio-temporal events pacing human preimplantation development. **a-b**, Streamgraph of gene module expression along pseudotime states. Each module was named by one of its representative genes. Expression level of each module is represented by the thickness of its ribbon on the streamgraph and corresponds to the WGCNA eigengene metric (**a**). Subdivision of the branches of the pseudotime model according to module expression: Pre-specification branch is subdivided into 3 states (1. Pre-morula, 2. Morula, 3. Unspecified B1/B2) and the TE branch is subdivided into 3 states (5. Early-TE, 8.TE.NR2F2-, 9.TE.NR2F2+). This yields a total of 9 states, numbered by their order of apparition in pseudotime (**b**). **c**, Projection of gene expression levels of CDH1 and TJP3 on the pseudotime model. **d**, Immunofluorescence of CDH1 (cyan), ZO1 (purple), Actin (yellow) and nuclear counterstaining (white) in indicated stages. **e**, Localization of CDH1 and ZO1 in a B4 blastocyst, showing ZO1 localized apically and CDH1 localized laterally. **f**, Projection of AQP3 gene expression levels on the pseudotime model. **g**, Immunofluorescence of AQP3 (yellow) and DAPI (white) as nuclear counterstaining, at indicated stages. Split channels and additional IFs are presented in **Extended Data Fig. 9**. Scale bars = 47 µm.

The gene modules identified here define the wave of genes, which pattern stages during preimplantation development, offering an opportunity to scrutinize the underlying molecular mechanisms involved. Indeed, during mouse preimplantation development, morphogenesis is immediately followed by transcriptomic specification (**Extended Data Fig. 1d**) whereas during human preimplantation development, transcriptomic specification occurs after cavitation (**Fig. 1f**). Studies in mammalians showed establishment of tight and adherent junctions at the morula stage, sealing the embryo^17-19^. Transcriptomic profile of adherent junction gene CDH1 and TJP3 tight junction protein in human are similar to mouse and bovine^17-19^ (**Fig. 2c**). Immunostaining of ZO1 and CDH1 during human preimplantation development confirmed basolateral expression of CDH1 from the morula stage, with subsequent apical recruitment of ZO1 (**Fig. 2d-e**). TJP1 is lowly expressed, which is not consistent with immunostaining of its related protein ZO1, but consistent with the expression profile of a closely related gene, TJP3. Another important event in the mammalian preimplantation development is the expression of aquaporins, at the morula/early blastocyst stage^19^. Remarkably, AQP3 gene is the only water channel expressed during human preimplantation development (**Fig. 2f**). Immunostaining for AQP3 revealed cytoplasmic membrane-localization at the morula stage, before being restricted apically upon sealing of the embryo by tight junctions (**Fig. 2g**).

Using our transcriptomic model, we sought markers of mature EPI and mature TE in B5 blastocyst, as this stage is easily characterized by hatching. Among transcription factors, we identified IFI16 as a candidate for mature EPI. B5 embryos displayed IFI16 restricted into ICM cells (**Fig. 3a**). Analysis of time-lapse assessed developmental stages showed no IFI16 expression before B2 and a robust expression after B4 (**Fig. 3b**). IFI16 is therefore specific to EPI, marking the onset of EPI proper. Identifying regulators of IFI16 could solve pathways involved in EPI specification and maturation. We also identified NR2F2, a transcription factor expressed in the TE branch after expansion. In B5 blastocysts, immunofluorescence showed nuclear localization of NR2F2 in TE, but restricted to the polar TE cells juxtaposed to the ICM marked with NANOG (EPI) (**Fig. 3c**). To investigate if NR2F2 is expressed readily in specified TE, we investigated B4 blastocyst, a stage immediately following lineage specification. Immunostaining revealed GATA3 and GATA2 expression throughout TE while NR2F2 was only expressed in the polar TE cells (**Fig. 3d**). Therefore, from E6 on, TE cells can be classified as NR2F2-positive and NR2F2-negative cells. NR2F2-positive cells localize at the end of the TE branch, the most mature part of TE signature. This supports a mechanism of TE maturation driven by contact with the ICM. Polar initiation of human TE maturation is consistent with the observation that the majority of human blastocysts attached to endometrial cells by the polar side with subsequent spreading of TE maturation to mural cells^20-27^. Analysis of most enriched signaling pathways pointed out prime candidates potentially driving maturation of EPI and TE: TGFβ, IGF1, BMP2, IL6 and FGF4 (**Extended Data Fig. 10**). Polar TE-driven maturation in human is reminiscent of the ICM-polar TE molecular dialog observed in mouse embryoids^17, 28^. However, human specific events might be involved as some genes critical for mouse TE maturation are not expressed in human, such as KLF6.

**Figure 3.**
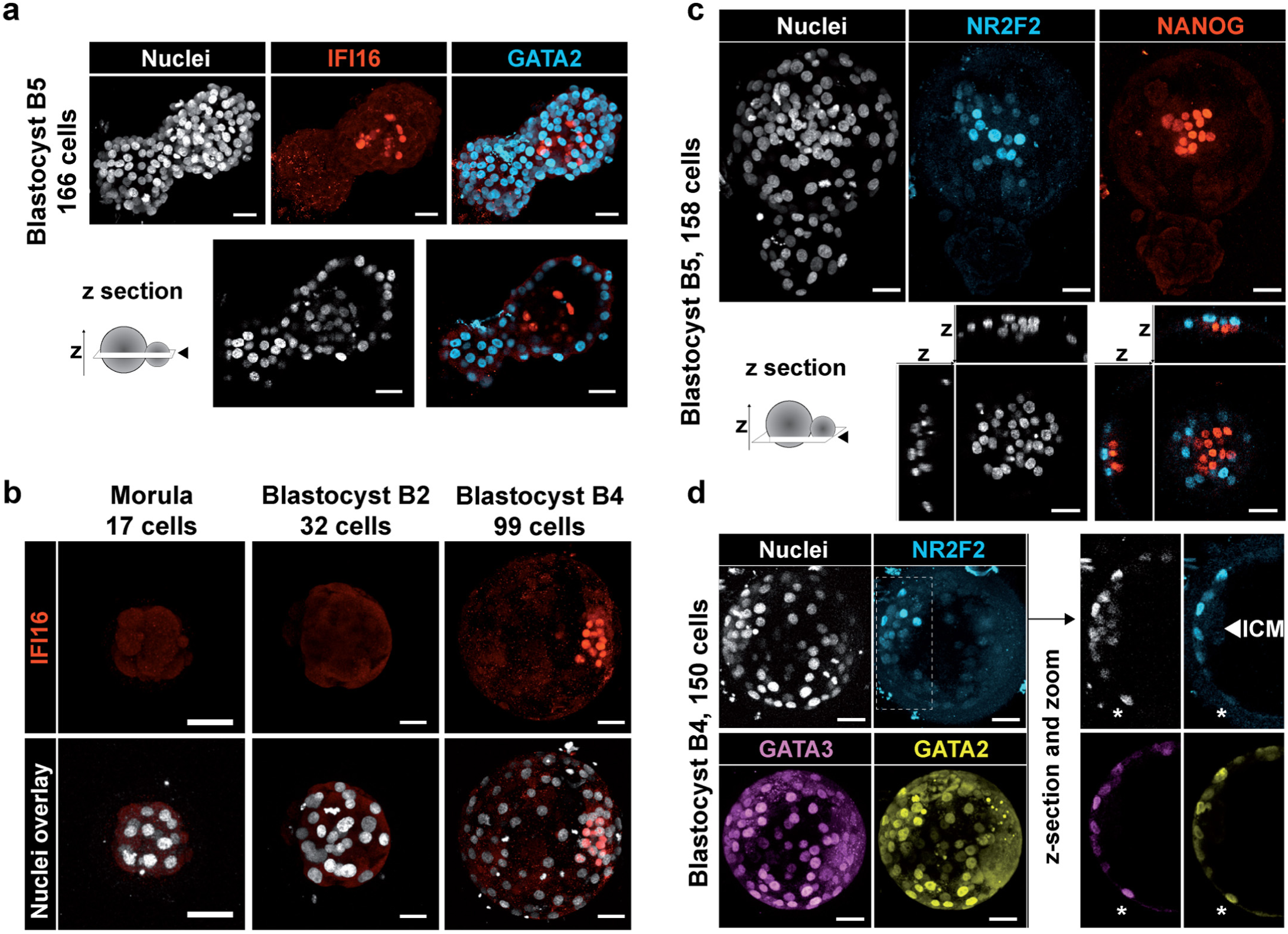
Identification of mature EPI (IFI16) and TE (NR2F2) markers. **a-b**, Immunofluorescence of IFI16 (red) co-stained with DAPI (white) nuclear counterstaining and GATA2 (cyan)(**a**) at indicated stages. *z-section* indicates a z-cutting plane. **c-d**, Immunofluorescence of NR2F2 (cyan) co-stained with DAPI nuclear counterstaining (white) and with NANOG (red)(**c**) at B5 stage or with GATA2 (yellow) and GATA3 (purple) at B4 stage (**d**). Asterisk marks a cell negative for NR2F2 and positive for GATA3 / GATA2. Scale bar = 47 µm.

Altogether, our study refines our understanding of the spatio-temporal events occurring during human lineage specification (**Fig. 4**). Our data suggests the EPI fate to be the first to be achieved, highlighted by genes expressed in all cells before specification then restricted to the EPI. This in turn triggers an EPI specific signature. TE is specified at B2/B3 with cells progressively losing expression of pluripotency-associated genes, correlating with loss of plasticity of TE cells^16^. The toughest lineage to resolve is PE, regardless of approaches undertaken^1, 3, 29^. A putative hypothesis is that PE is not fully established during human preimplantation development.

**Figure 4.**
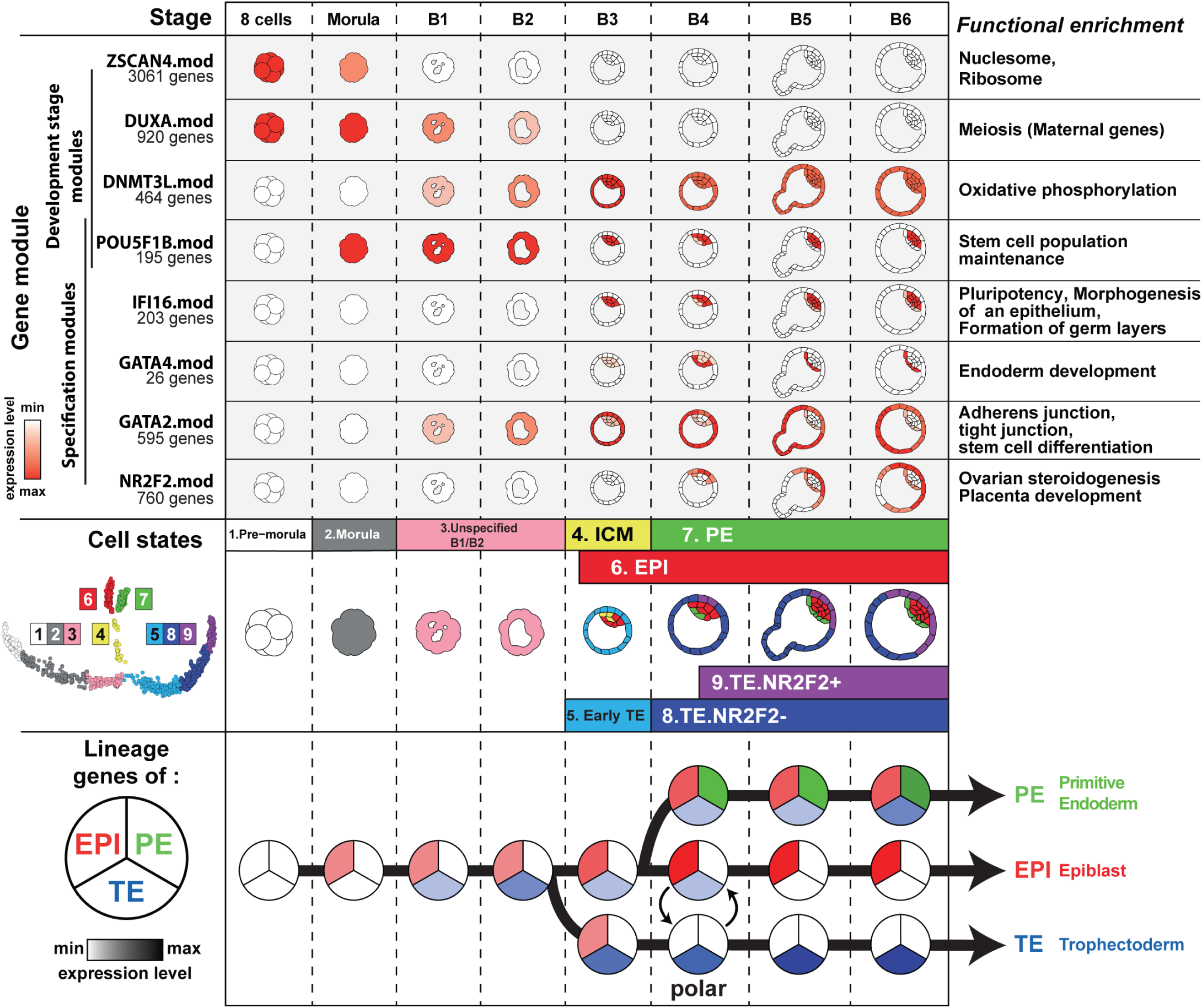
Human preimplantation development model. Each annotation is represented per morphology-based stages of human preimplantation development. Upper panel: Schematic representation of gene module expression. Functional enriched functions are presented on the right. Complete enrichment data are presented in **Extended Data Fig. 8**. **b.** Middle panel: Schematic of pseudotime cell states spatiotemporal localization. A projection of cell states on the pseudotime model is shown on the left. Lower panel: expression of lineage specific stage per stage and per cell fate.

Our study highlighted signaling pathways and events involved during human preimplantation development linked to IVF. Our model therefore paves the way to more efficient media formulation, and readouts to assess development of human embryos. Improvement of IVF procedures is necessary since the procedural average efficiency is below 27%^30^. Our study contributes to an improved understanding of human preimplantation, a gateway to improve IVF success rates.

## Extended Data Information line

There are 10 Extended Data figures and 4 Extended Data Tables.

### Acknowledgements

We thank our colleagues F. Lanner and K. Niakan for sharing data and providing feedback. We thank J. Jullien, V. Pasque and J. Chappel for critical review of the manuscript. DM is supported by FINOX and MSD contributed to the project. We thank core facilities: BIRD, PFiPSC and MicroPicell. This work was supported by “Paris Scientifique region Pays de la Loire: HUMPLURI” and IHU CESTI.

## Author contributions

TF and LD designed the study. DM, SL and LD wrote the manuscript with input from all authors. SL, AR, JF, JL performed embryo manipulation. PH, SN, SL, AR and LD performed IF analysis. MS and TM performed single-cell RNAseq. DM, VFC, YL, performed bioinformatics analysis under supervision of AB and JB and input from BB and GC. TF and PB supervised human embryo donation. All authors approved the final version of the manuscript.

## Author information

All original data have been deposited on XXX under number XXX.

Competing interest declaration: D.M. is supported by FINOX Biotech FORWARD initiative 2016.

## Methods

### Human preimplantation embryos

The use of human embryo donated to research as surplus of IVF treatment was allowed by the French embryo research oversight committee: Agence de la Biomédecine, under approval number RE13-010. All human preimplantation embryos used in this study were obtained from and cultured at the Assisted Reproductive Technology unit of the University Hospital of Nantes, France, which are authorized to collect embryos for research under approval number AG110126AMP of the Agence de la Biomédecine. Embryos used were initially created in the context of an assisted reproductive cycle with a clear reproductive aim and then voluntarily donated for research once the patients have fulfilled their reproductive needs or tested positive for the presence of monogenic diseases. Informed written consent was obtained from both parents of all couples that donated spare embryos following IVF treatment. Before giving consent, people donating embryos were provided with all of the necessary information about the research project and opportunity to receive counselling. No financial inducements are offered for donation. Molecular analysis of the embryos was performed in compliance with the embryo research oversight committee and The International Society for Stem Cell Research (ISSCR) guidelines^31^.

### Human preimplantation embryos culture

Human embryos were thawed following the manufacturer’s instructions (Cook Medical: Sydney IVF Thawing kit for slow freezing and Vitrolife: RapidWarmCleave or RapidWarmBlast for vitrification). Human embryo frozen at 8-cell stage were loaded in a 12-well dish (Vitrolife: Embryoslide Ibidi) with non-sequential culture media (Vitrolife G2 plus) under mineral oil (Origio: Liquid Paraffin), at 37°C, in 5% O_2_/6% CO_2_.

### Human embryo time-lapse imaging

Embryos were loaded into the Embryoscope® (Vitrolife®), a tri-gas incubator with a built-in microscope allowing time-lapse monitoring of embryo development. Images were captured on seven focal plans every 15-min intervals using Hoffman modulation contrast (HMC) optical setup^1^ and a 635 nm LED as light source as provided in the Embryoscope®. The resolution of the camera is 1280×1024 pixels. The development of each embryo was prospectively annotated as described by Ciray et al., by two trained embryologists undergoing regular internal quality control in order to keep inter-operator variability as low as possible^32^. Zona Pellucida (ZP) thickness was measured by our novel analysis pipeline (*Feyeux, Reignier et al, biorxiv 2018*). The term tM refers to a fully compacted morula. At the blastocyst stage, tSB is used to describe the onset of a cavity formation, tB is used for full blastocyst i.e the last frame before the Zona Pellucida (ZP) starts to thin, tEB for expanded blastocyst, i.e. when the ZP is 50% thinned. Blastocyst contractions and the beginning of herniation were also recorded.

### Immunofluorescence of human embryos

Embryos were fixed at the morula, B1, B2, B3, B4, B5 or B6 stages according to the grading system proposed by Gardner and Schoolcraft^33^. Embryos were fixed with 4% paraformaldehyde for 5 min at room temperature and washed in PBS/BSA. Embryos were permeabilized and blocked in IF Buffer (PBS–0.2% Triton, 10% FBS) at room temperature for 60 min. Samples were incubated with primary antibodies over-night at 4°C. Incubation with secondary antibodies was performed for 2 hours at room temperature along with DAPI counterstaining. Primary and secondary antibodies with dilutions used in this study are listed in **Extended Data Table 1**.

### Imaging

Confocal immunofluorescence pictures were taken with a Nikon confocal microscope and a 20× Mim objective. Optical sections of 1µm-thick were collected. The images were processed using Fiji (http://fiji.sc) and Volocity visualization softwares. Volocity software was used to detect and count nuclei.

### Single-cell RNA sequencing

Single-cell isolation and overall dataset are presented in **Extended Data Fig. 2**. Single-cell RNA-seq libraries were prepared according to the SmartSeq2 protocol^34, 35^ with some modifications^36^. Briefly, total RNA was purified using RNA-SPRI beads. Poly(A)+ mRNA was reverse-transcribed to cDNA which was then amplified. cDNA was subject to transposon-based fragmentation that used dual-indexing to barcode each fragment of each converted transcript with a combination of barcodes specific to each sample. In the case of single cell sequencing, each cell was given its own combination of barcodes. Barcoded cDNA fragments were then pooled prior to sequencing. Sequencing was carried out as paired end 2×25bp with an additional 8 cycles for each index. The FASTQ files were mapped with Hisat2^37^ on GRChH38 genome version, downloaded from ensembl.org. HTSeq41 was used to generate raw counts tables from BAM files, using the matching GTF for the reference genome.

### Raw count table treatments

Samples were filtered with the use of the R function *isOutlier* from SCRAN^11^ library. This function tags samples as outliers with a threshold based on median derivation away from the median of the metric. Samples were filtered with two metrics: the number of expressed genes with a threshold of 2 median away derivation from median, and the total number of counts in the sample with a threshold of 3 median away derivation from median. Both metrics were used to discard sample in a two-sided way, below and above the median. Genes that were expressed in less than two cells and with an average expression less than 0.1 counts were removed.

The four datasets were then normalized together using the *computeSumFactor* function from SCRAN. Logged and non-logged data were collected using the *normalize* function from scater R library. To compute batch effect free expression, we normalized the data as described above but per dataset. We used mutual nearest neighbors correction implemented by the function *mnnCorrect* to achieve the batch correction from the log-normalized data. The reference dataset that were used for *mnnCorrect* is from Petropoulos *et al*^*1*^.

### Computation of pseudotime model

Monocle2^8^ needs a subset of gene to make the dimensionality reduction and pseudotimes trajectories. To choose the best set of ordering genes we took samples that passed the quality control from Petropoulos et al.^*1*^ to avoid batch effects. We used SCRAN for processing size factors (normalization factor). We then created an R object with the *newCellDataSet* object used by Monocle2 with the raw expression and the expressionFamily parameter set as “negbinomial.size()”. Size factors were attributed according to the SCRAN results. The next step consisted of estimating empirical dispersion of each gene in the negative binomial model with the *estimateDispersions* function. We used the dispersion table function to gather the empirical dispersion and the fitted theoretical dispersion for each gene. We made a ratio of empirical dispersion on the theoretical dispersion for each gene. This ratio describes an over-dispersion score.

For a given gene *i* the over-dispersion score S is calculated as follows: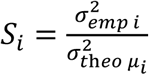

Genes with an average log expression < 0.5 across samples were filtered out. Remaining genes were ranked based on their overdispersion score.

Pseudotimes models were generated using a range of top ranked ordering genes from the top 500 to the top 5500. This lead to 5000 pseudotime models. For each pseudotime model, A new R object was created with the newCellDataSet function, with the batch corrected expression from the four datasets as input and the expressionFamily parameter set as “gaussianff()”. Selected ordering genes were then set as input of Monocle2 algorithm, with the number of resulting dimensions set to three dimensions. An automatic classification of pseudotime models was set up following three criteria based on their topology:

- Number of branches populated by mural TE cells.
- Succession of developmental stage.
- Position and number of branching points.

The most common topologies were: (i) all mural TE cells within one branch, (ii) developmental stages succeeding one other, i.e. morula between 8-cell and blastocysts, (iii) two branching points, (iv) first branching point at E5. The chosen pseudotime model belonged systematically to the most abundant topologies and is calculated from 4484 ordering genes. The resulting 3-dimension pseudotime model was rotated to obtain a 2-dimensional projection.

### WGCNA

WGCNA^14^ was performed on batch corrected data using a soft power of 10 with signed Pearson correlation. Resulting module were manually curated to choose a set of 8 modules that were well represented in data and that have distinct behaviors. For each module we use the module eigengene metric that is given by WGCNA to infer the global module expression across the samples. A loess regression of eigengene by pseudotime was used for **Fig. 2a**.

### Enrichment analysis

Module enrichment analysis was performed with FGSEA^38^. Ranking metric for FGSEA was set as WGCNA gene module membership. This score is processed by Pearson correlation of module eigengene and gene expression. Enrichment was made on five databases: Gene ontology (Cellular Component, Molecular Function and Biological Process), Reactom and KEGG. All retained term were enriched below an adjusted Benjamini-Hochberg p-value of 0.05. A list of transcriptional factors (TF) was downloaded from the RegNetwork database^39^. Pathway eigengene metric (**Extended Data Fig. 8**) was processed by taking the first component from a principal component analysis based on the genes from each enriched term.

### Loess regressed expression by pseudotime

We used a Locally Weighted Regression (LOESS) to fit expression in the Pseudotime by cell fate with neighbor impact of 0.75. Expression profiles of common segments were fitted to extract global tendencies. A last LOESS was computed with a low neighbor impact to merge segments to obtain continuous expression curves.

### Subdivision of pseudotime branches

The original pseudotime model was constituted by five states, as Monocle2 separates states only by branching point. We subdivided pre-specification and trophectoderm branch samples by using WGCNA module eigengene. For both branches, WGCNA modules with a Pearson correlation higher than 0.75 with the pseudotime were selected. A loess regression of these module eigengene by pseudotime was performed, followed by a hierarchical clustering of regressed module eigengenes. The clustering was then partitioned. For each branch the best partition was determined at three clusters with the greatest relative loss of inertia method.

### Data representation used in each figure

Raw expression:

- estimating gene dispersion and select ordering genes

Normalized expression:

- projection of gene expression on pseudotime (**Fig. 1c**, **Extended Data Fig. 1f**)
- Pseudotime User Interface

Loess regressed expression by pseudotime:

- heatmap (**Extended Data Fig. 7b**)
- expression profile curves (**Fig. 2c, Fig. 2f**, **Extended Data Fig. 7a**, **Extended Data Fig. 10a-d**)

Batch corrected logged expression:

- computation of pseudotime model
- WGCNA

ComplexHeatmap^40^, ggplot2 and d3.js. were used for graphical representation. Hierarchical clustering was done using the Ward criteria and from a correlation distance for the gene/pathway eigengenes, or from the euclidean distance for other metrics.

### Pseudotime User Interface (PTUI)

PTUI was developed using HTML5 and d3.js.

### Mouse single-cell RNA-Seq analysis

Mouse dataset were analyzed in a similar way to human datasets, without batch correction. Alignment step was done from the mm10 version of the genome. Timing of blastulation is corroborated by time-lapse (*Feyeux, Reignier et al, biorxiv 2018*).

### RNA velocity

RNA velocity was performed from BAM of samples that have passed all quality control in the final counts table. First, we used velocyto.py^41^ using the command *velocyto run*, with the parameter --logic as “SmartSeq2”, and the parameter -m (RepeatMasker annotations) as a GTF downloaded from the UCSC genome browser. The global GTF was the same that were used for the computation of raw counts table. Resulting loom files were merged using loompy.combine from lompy python package. We used velocyto.R for computing Velocity matrix. Loom files were read with the function *read.loom.matrices*. Then we separated spliced reads matrix, unspliced reads matrix and spanning reads matrix. For each of the matrix a gene filtering was done with the function *filter.genes.by.cluster.expression*. The min.max.cluster.average parameters were set for the corresponding matrix as:

- spliced reads matrix: 5
- unspliced reads matrix: 1
- spanning reads matrix: 0.5

Then RNA velocity was estimated using *gene.relative.velocity.estimates*, with the following parameters: fit.quantile = 0.05, deltaT = 1, kCells = 5.

PCA of **Fig. 1g-h** were calculated with the function *pca.velocity.plot*.

### Data and software availability

Single cell alignment pipeline: *available soon*

WGCNA: *available soon*

Pseudotime User Interface source code: *available soon*

All other parts of the code are available upon request.

## Extended Data Table Legends

**Extended Data Figure 1.**
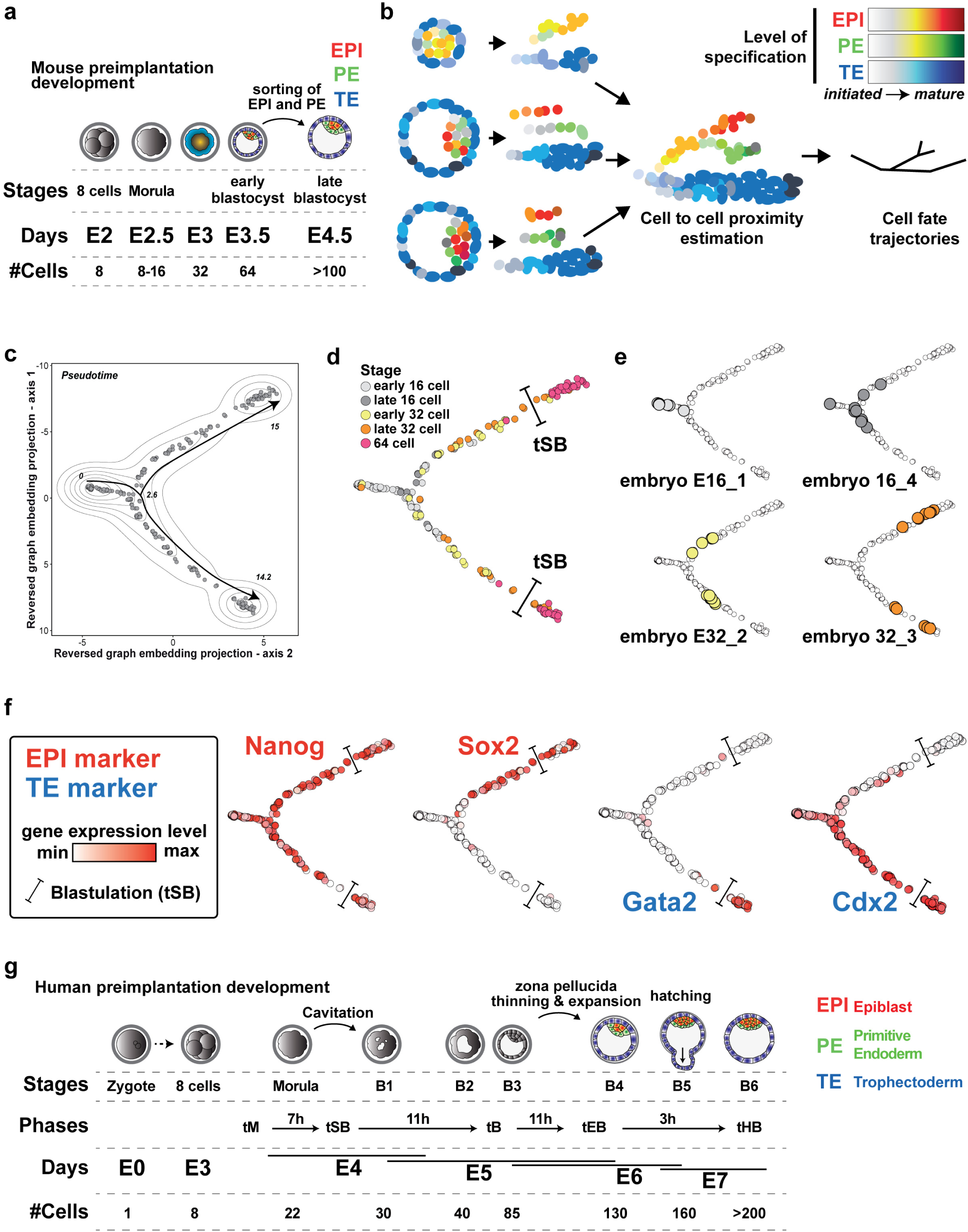
Mouse preimplantation pseudo-time model. **a**, Schematic of mouse preimplantation development. **b**, Schematic presentation of pseudotime analysis principle. All single-cell transcriptomic profiles are combined to create the likeliest probabilistic path from one cell to another. **c-e**, Result of mouse preimplantation development pseudotime analysis, based on the dataset of Posfai et al. Model with cell density and pseudotime values (**c**). Developmental stages are projected on the pseudotime model (**d**), with 4 particular embryos highlighted (**e**). tSB means blastulation, the onset of the blastocoele cavity. **f**, Projection of expression levels of 4 indicated genes. **g**, Schematics highlighting staging differences between mouse and human. In IVF clinics, “Stages” and “Phases” are interchangeable. Average time scale for fresh embryos are shown between phases: morula compaction (tM), starting of blastulation (tSB), blastocyst stage (tB), initiation of expansion of the blastocyst (tEB) and hatched blastocyst (tHB). Number of cells indicated is an average from what we observed and from published IF^24^.

**Extended Data Figure 2.**
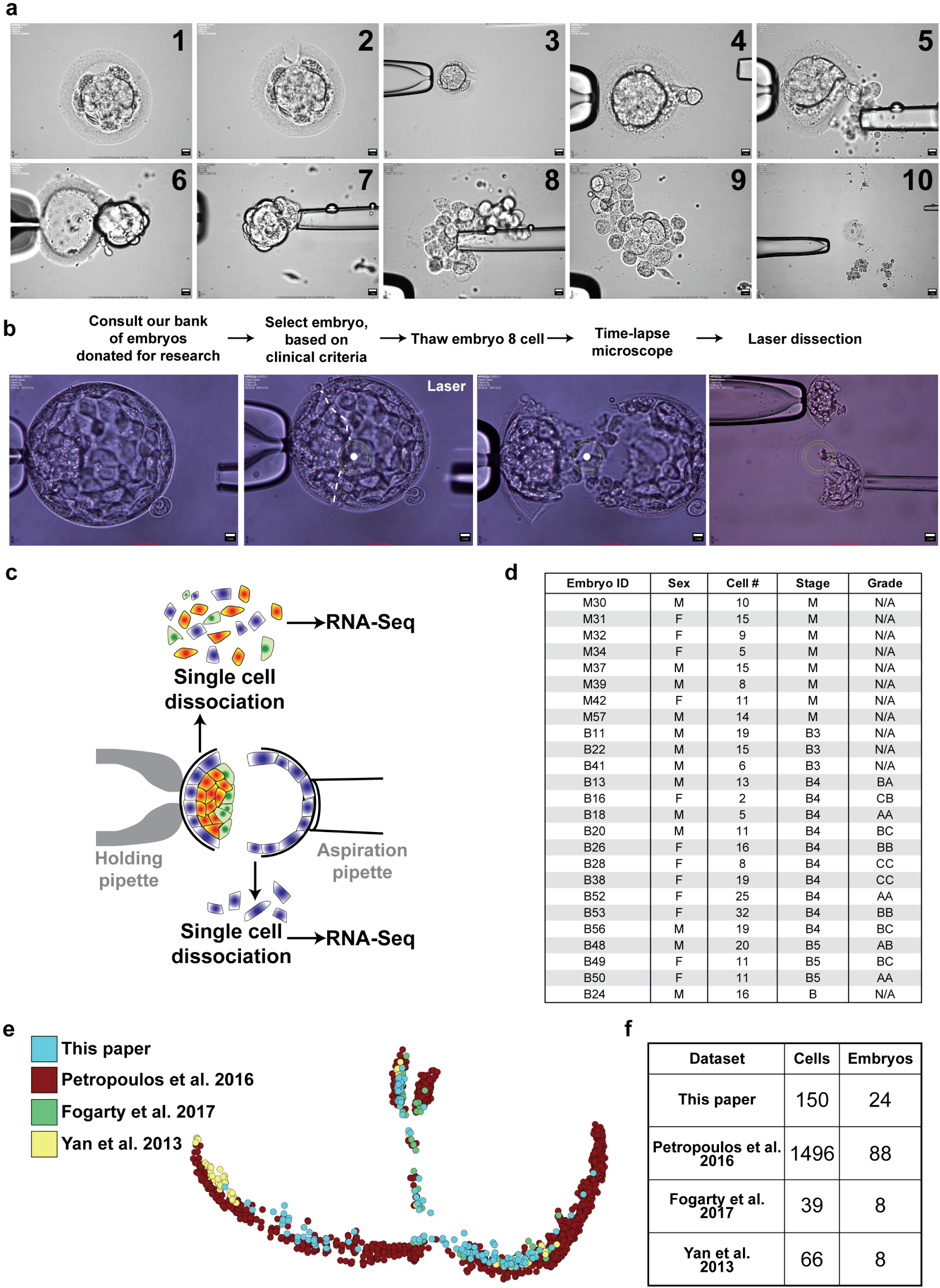
Sample preparation and dataset overview. **a-c**, Detailed manipulation of embryos before scRNAseq. Morula are first incubated in decompaction media, then the zona pellucida (ZP) is pierced by a laser (dot). Using micropipettes, fragments are removed, and the morula is extracted from the ZP. Cells are then manipulated until dissociated (**a**). For blastocysts, embryos are thawed and monitored until desired developmental stage (**b**). Embryos are then laser-dissected, the polar side (containing TE, PE and EPI cells) and the mural side (containing TE cells) (**c**). **d**, Detailed sampling of our dataset. For each embryo, the embryo ID, sex, number of cells sequenced, stage and grade are indicated. **e-f**, Projection of samples per dataset on the pseudotime model (**e**) and sampling of the four datasets (**f**).

**Extended Data Figure 3.**
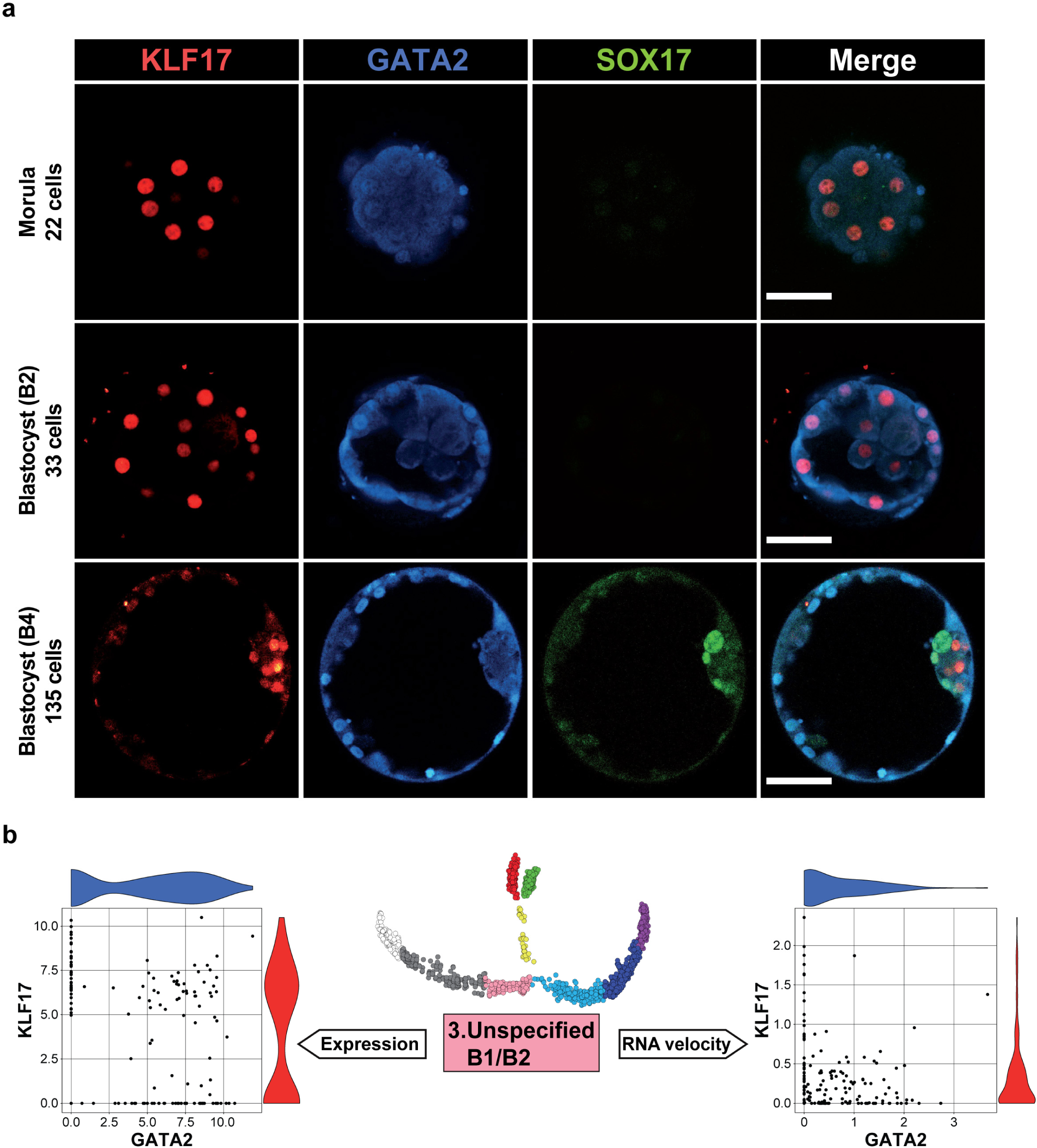
Protein expression, RNA expression and RNA velocity of GATA2 and KLF17. **a**, Immunofluorescence for KLF17 (red), GATA2 (blue) and SOX17 (green) for morula, B2 and B4 Blastocyst. Scale bar = 50 µm. Figure from Kilens et al. (2018). **b**, RNA expression and RNA velocity for KLF17 and GATA2 for cells from the pre-specification state.

**Extended Data Figure 4.**
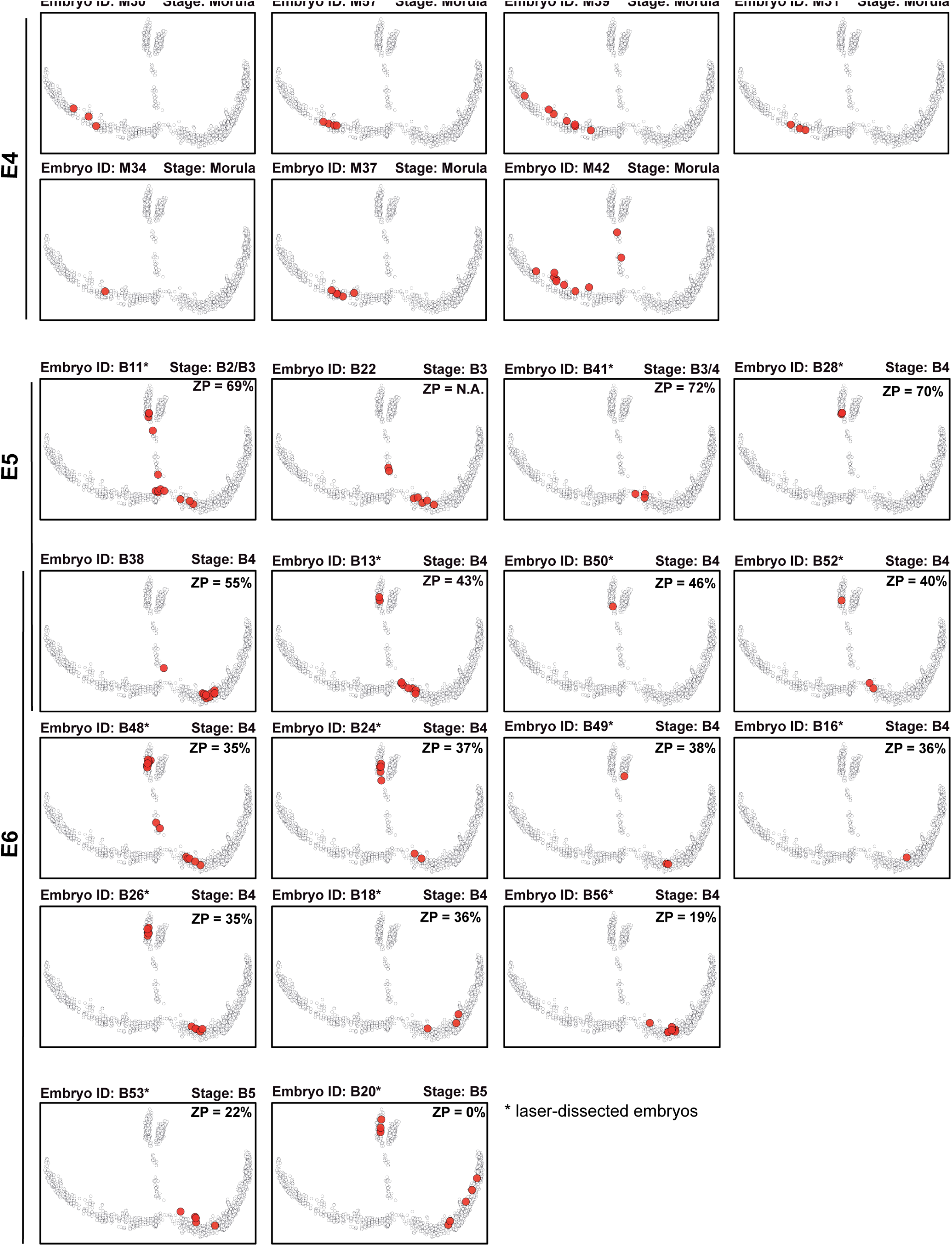
Projection of samples per embryo on the pseudotime model. All cells from each embryo sequenced for this paper are projected on the pseudotime model. The stage of the embryo has been annotated by time-lapse microscopy and the zona pellucida thickness has been measured and is indicated as a percentage of original size. See Feyeux et al (biorxiv) for more details.

**Extended Data Figure 5.**
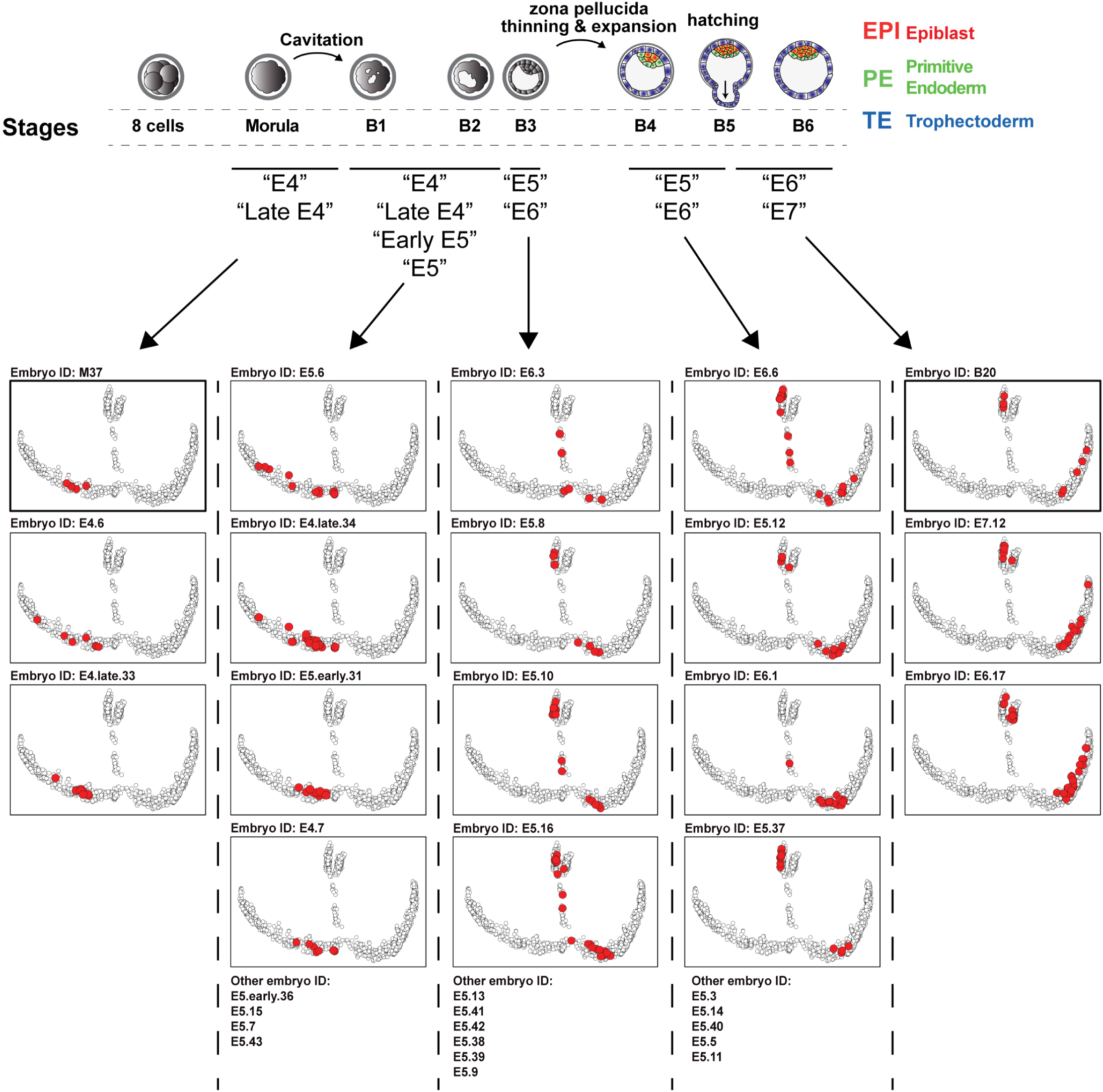
Refined annotation from all cells and embryos included in our study. Embryos are ranked based on the average positioning of all cells from each embryo included in the study. This highlights that embryos annotated “E5” can be found at the exit of morula, just before specification, just after specification or later on, interspersed with “E6”. This could be due to difficulty to annotate embryos based on a single observation, on the fact that multiple morphologies are present at day 5, or simply because the subset of cells sequenced from an embryo is not representative of the molecular state of the embryo. Our model allows us to bin embryos based on their molecular profile, allowing a finer analysis of events pacing human preimplantation development, especially at E5.

**Extended Data Figure 6.**
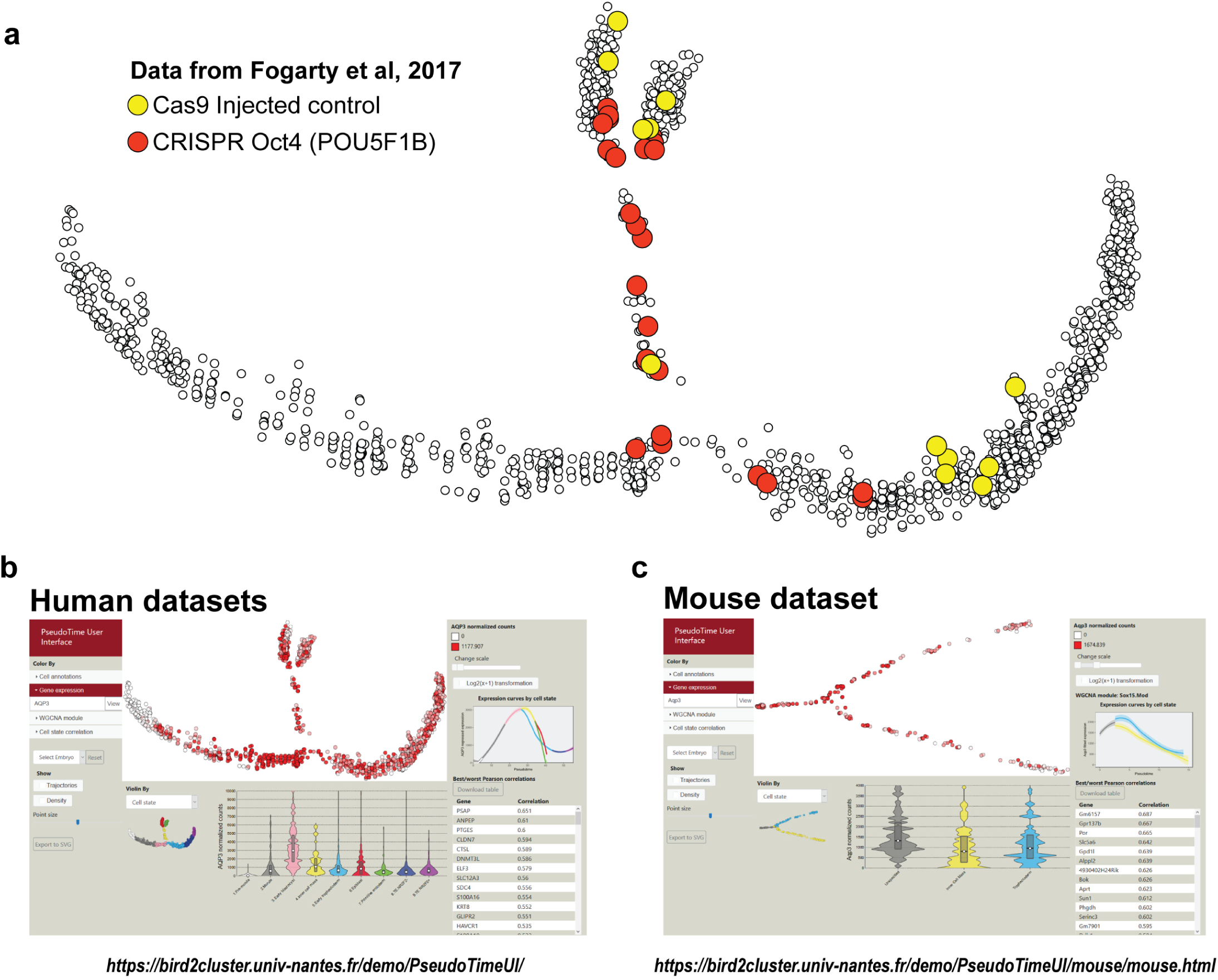
Projection of KO embryos for OCT4 on the pseudotime and overview of the pseudotime user interfaces (PTUI). **a**, Projection of human single-cells KO for POU5F1 (orange) or controls (yellow). Samples from Fogarty *et al.* 2018. **b-c**, Screenshot of our human (**b**) and mouse (**c**) web pseudotime user interface. URLs are indicated below.

**Extended Data Figure 7.**
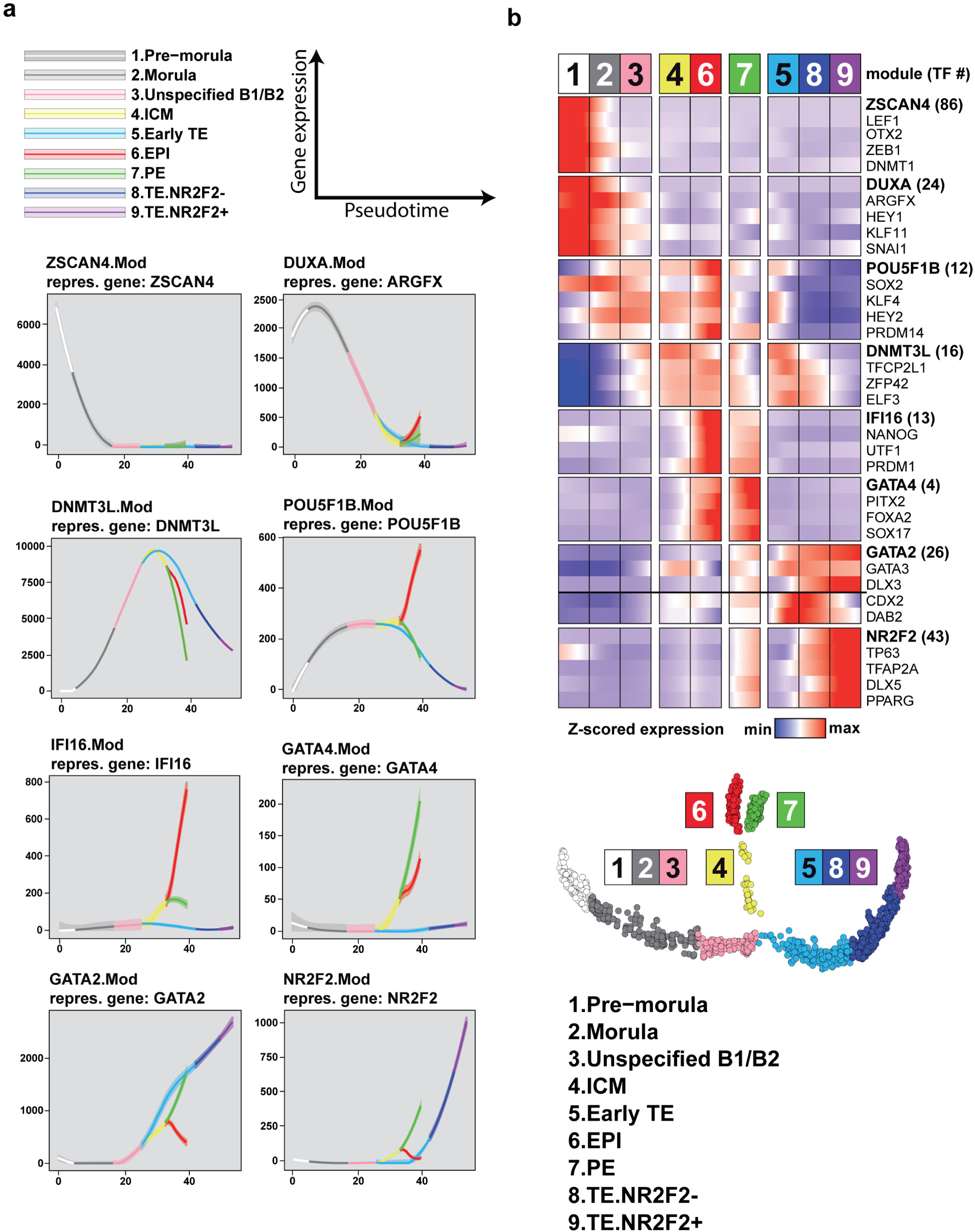
Tendency curves for WGCNA modules and functional enrichment analysis. **a**, Expression profile of a representative gene of each gene module. **b**, Expression levels of transcription factors specific of gene modules. The heatmap is subdivided by states and ordered by pseudotime for each state. Genes were selected by their membership degree for the gene module.

**Extended Data Figure 8.**
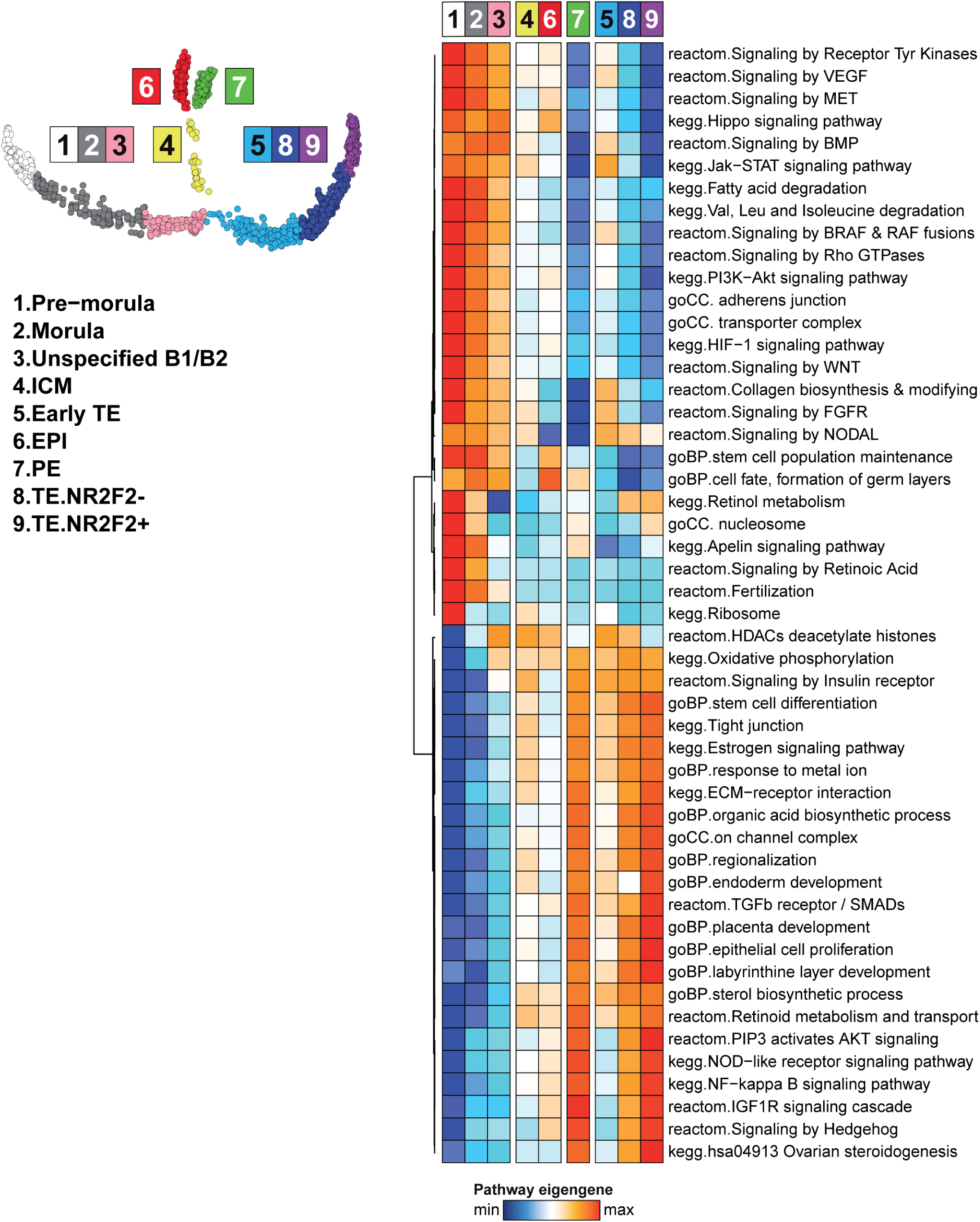
Functional enrichment per pseudotime states. Heatmap representing the average of 51 pathway eigengene across the pseudotime states. Pathway eigengene is defined as the first principal component of a principal component analysis with the genes of the pathway as initial dimensions. Enrichment was done in gene modules according to 5 databases: Gene ontology (GO Molecular Function, GO Cellular component, GO Biological process), KEGG and Reactom. All terms were significantly enriched in at least one gene module with an adjusted pvalue < 0.05.

**Extended Data Figure 9.**
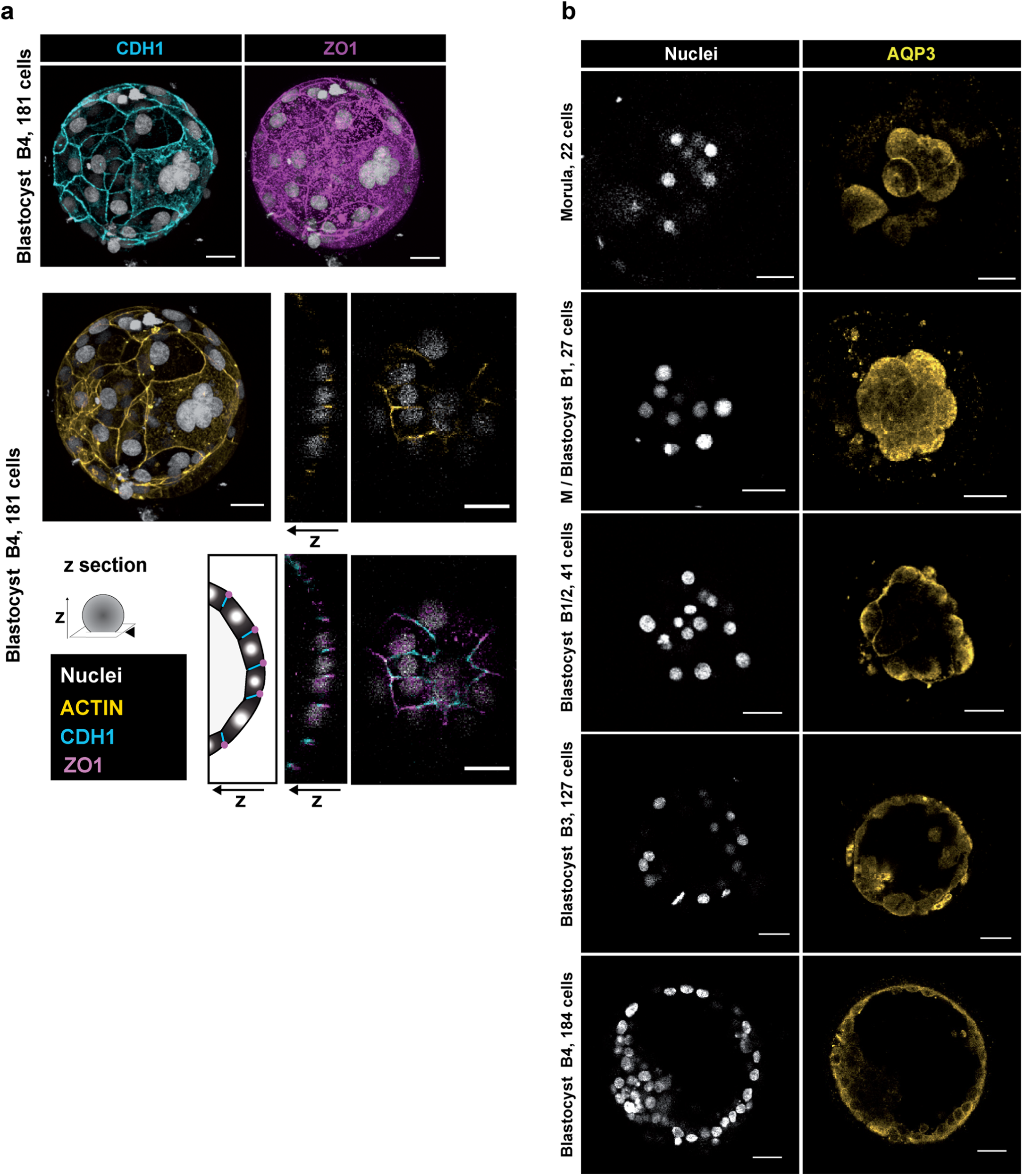
Additional immunofluorescence visualization. **a**, Extended focus visualization of immunofluorescence of ACTIN (yellow), CDH1 (cyan) and ZO1 (purple) with nuclear counterstaining (white) for a B4 blastocyst. **b**, Split-channels immunofluorescence for AQP3 (yellow) with nuclear counterstaining (white) at indicated stages. Scale bar = 47 µm.

**Extended Data Figure 10.**
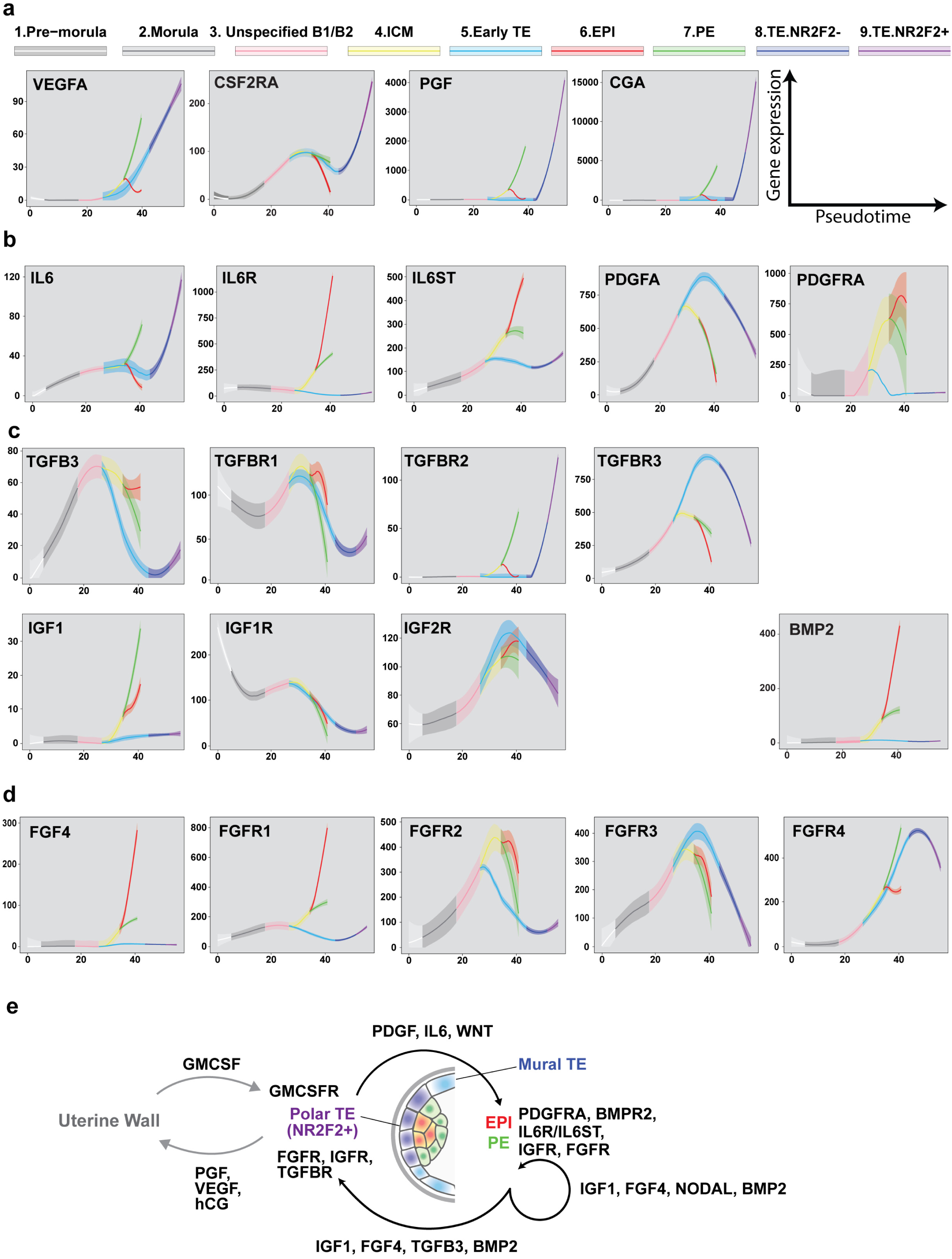
Expression profile of selected cytokines and receptors. **a-d**, Expression profiles signaling pathways components during preimplantation development. Cytokines secreted by the TE targeting the uterine wall (**a**), cytokines expressed by the TE and targeting the ICM (**b**), cytokines expressed by the ICM and targeting the TE (**c**) and cytokines expressed by the ICM targeting the whole embryo (**d**). **e**, Schematic representing cytokine-receptors loops identified in enriched pathways.

**Extended Data Table 1 | Antibodies**

Detail of primary and secondary antibodies used for this study.

**Extended Data Table 2 | Cells annotations for human dataset**

Annotation for each cell used in the study of human preimplantation development. Annotations beginning with “Author” are referring to annotation of author’s dataset. “Author.lineage”: lineage attribution of authors. “Author.TEside”: Trophectoderm side attribution from Petropoulos et al. (2016). “Author.pseudoTime”: pseudotime of each cell according to Petropoulos et al. (2016). “StirparotEtAl.lineage”: lineage attribution of dataset from Petropoulos et al. (2016) by Stirparo et al. (2018).

**Extended Data Table 3 | Cells annotations for mouse dataset**

Annotation for each cell used in the study of human preimplantation development. Annotations beginning with “Author.lineage” is referring to the lineage attribution of Posfai et al. (2017).

**Extended Data Table 4| Gene modules**

Detailed gene list for each WGCNA modules.

